# Making sense of RNA-Seq data: from low-level processing to functional analysis

**DOI:** 10.1101/010488

**Authors:** Oleg V. Moskvin, Sean McIlwain, Irene M. Ong

## Abstract

Numerous methods of RNA-Seq data analysis have been developed, and there are more under active development. In this paper, our focus is on evaluating the impact of each processing stage; from pre-processing of sequencing reads to alignment/counting to count normalization to differential expression testing to downstream functional analysis, on the inferred functional pattern of biological response. We assess the impact of 6,912 combinations of technical and biological factors on the resulting signature of transcriptomic functional response. Given the absence of the ground truth, we use two complementary evaluation criteria: a) consistency of the functional patterns identified in two similar comparisons, namely effects of a naturally-toxic medium and a medium with artificially reconstituted toxicity, and b) consistency of results in RNA-Seq and microarray versions of the same study. Our results show that despite high variability at the low-level processing stage (read pre-processing, alignment and counting) and the differential expression calling stage, their impact on the inferred pattern of biological response was surprisingly low; they were instead overshadowed by the choice of the functional enrichment method. The latter have an impact comparable in magnitude to the impact of biological factors *per se*.

## Introduction

The ultimate goal of any transcriptomic experiment is to discover functional patterns of biological response to conditions of interest (treatments, environmental influences, mutations etc). However, transcriptomic analytical pipelines consist of multiple processing stages, and one would reason that the propagation of results from analytical choices made at earlier stages to the downstream steps might lead to vastly different conclusions. To the best of our knowledge, systematic comparison of the results between different combinations of analytical methods within a pipeline in context of the resulting functional signature of cellular response has not been previously addressed.

Soon after its invention, RNA-Seq^1^ started to aggressively replace microarrays in the field of transcriptomics^2^, and this shift is due to RNA-Seq’s impressively larger (at least 2 orders of magnitude) dynamic range and its ability to detect novel transcripts^1^.

Both microarray and RNA-Seq technologies have their specific biases: given the same concentration of target cDNA, number of probe-target duplexes on a microarray will be strongly sequence-dependent^3^ and even physical position of the probe on the array may have an influence on the measurement by “fogging” the positions of low-intensity probes with the fluorescence emitted from their bright neighbors, resulting in artificial increase in estimated expression of numerous genes^4^. The observed phenomenon of “probe affinity effects” was addressed in low-level microarray processing algorithms such as RMA^5^. Recently, the phenomenon appeared to be in the focus of more mechanistic modeling efforts^6^. On the other hand, RNA-Seq has its own biases, starting from gene length bias that could be eliminated by normalizing for RNA length and for the total number of mapped reads^7^ and as uniform sampling of mRNA pool ^8^ to biases that belong to next-generation sequencing technology itself such as nucleotide per cycle bias and mappability bias^9^.

The nature and associated problems of hybridization- and sequencing-based transcriptomic technologies led to believe that microarrays and RNA-Seq will stay complementary^10^; this was supported by co-existence of reports that show RNA-Seq to be either more sensitive than microarrays^11^ or surprisingly less sensitive for low-expressed genes^12^. Apparently, the difference in conclusions stems directly from the lack of consistent, well-established data processing routines. Long-term, one may expect prospering of RNA-Seq based on its objective advantages of high dynamic range and *de novo* transcriptome structure discovery, while all the technical issues are addressed by the analytical methods. However at present, the methods of RNA-Seq analysis are still rapidly evolving, and most importantly, detection of biologically meaningful responses remains a “bigger challenge” within both microarray and RNA-Seq worlds^13^. This makes investigating the interface between data processing stages and functional pattern detection essential.

We use two orthogonal approaches to evaluate the results of various combinations of RNA-Seq pipeline processing: gene-level consistency of DE results on microarray and RNA-Seq platforms and – the key of this study - conservation of functional signatures in 2 biologically related comparisons within the RNA-Seq platform.

The published results by Schwalbach^14^ and Keating^15^ that compare three different conditions of E.coli growth: control (SynH, synthetic corn stover hydrolysate without the toxic components), naturally toxic medium (ACSH, corn stover hydrolysate containing toxic components of lignin decomposition) and artificially toxic medium (SynHLT, synthetic corn stover hydrolysate with added mixture of toxic components identified in ACSH) were used to help evaluate the relative impact of various stages of RNA-Seq processing on the inferred functional patterns of cellular response.

Our study shows that the lower level stages of the RNA-Seq analytical pipeline, while being highly diverse in nature and options, have surprisingly low influence on the derived functional pattern of the response. At the same time, choice of the functional enrichment methodology is found to be critical.

## Results

### Overview

Our approach to evaluating various methods within an RNA-Seq data analysis pipeline is outlined in Fig.1. For **e**very major stage of the pipeline (low-level processing, differential expression calling and functional pattern detection) we evaluate the results using both internal and external criteria whenever possible. The internal criteria are restricted to a particular processing stage and do not rely on external information, e.g. results from a different gene expression measuring platform, or biological knowledge. Every evaluation stage will be described in the respective subsection of the Results. The key external evaluation approach used in this study is correlation between vectors of inferred functionality for the two biologically related comparisons (Fig.2). The level of similarity of the functional profiles derived for the two comparisons was selected as an external biological criterion because: a) the two toxic media are expected to cause similar profiles of cellular response because the composition of the artificially toxic medium was carefully matched to the composition of the naturally toxic medium and b) this expectation was recently supported experimentally^15^. In order to check the robustness of higher-level functional pattern detection, we have chosen two comparisons that are diverse enough to depart from checking gene-level signal reproducibility and similar enough to expect similar higher-level patterns of response.

**Fig. 1.**
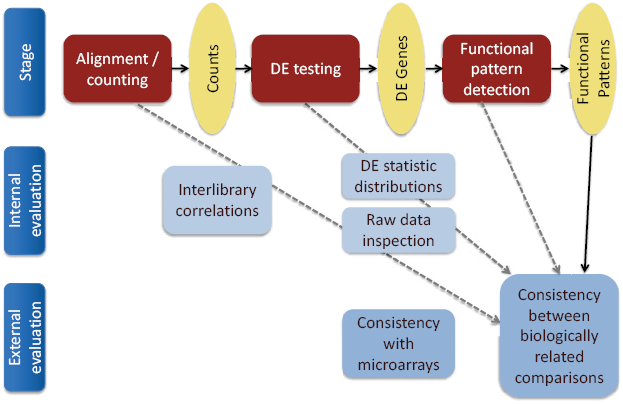
Overview of multilevel RNA-Seq pipeline evaluation approach.

**Fig. 2.**
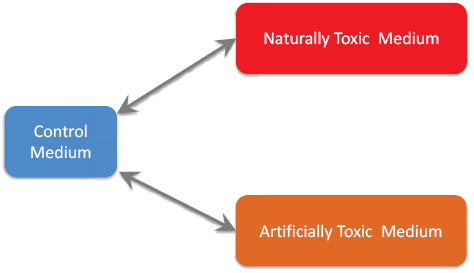
Related biological comparisons used. Naturally toxic medium: ammonia-pretreated corn stover hydrolysate (ACSH); artificially toxic medium: synthetic hydrolysate with added cocktail of toxic compounds discovered in ACSH (“lignotoxins”); control medium: synthetic hydrolysate with lignotoxins omitted. See ^15^ for further details.

In order to consider influences of both technical and biological nature on the same scale, the relative contributions of various processing stages were assessed in comparison with contribution from biological factors. For this purpose, all the combinations of the various methods were used to analyze time series data consisting of 3 growth stages (exponential, transitional and stationary^15^).

Exhaustive combining of 2 read pre-processing strategies, 2 alignment / counting pipelines, 3 count normalization methods, 2 pairwise comparisons, 4 DE calling methods, 3 functional overrepresentation strategies, 4 geneset types, 2 directions of expression changes, and 3 time points resulted in 6,912 vectors of functional enrichment results (approximately 1 million of individual gene set enrichment p-values). After removal of vectors with NA values, the pool of 3,145 pairwise correlations between the vectors representing the two biologically related comparisons was modeled as a function of all the technical processing factors plus biological factor (growth stage). This provided a big picture estimate of factor influence which complements the results of stage-specific evaluations.

### Low-level processing and alignment / counting

We considered two options for read pre-processing: “RAW” (untrimmed) vs. “QC” (trimmed reads) in order to investigate the tradeoff between having more sequence data and having more confident base calls at every base position in a read. For alignment/counting, we considered “BWA-HTSeq” vs. “Bowtie-RSEM” (see Materials and Methods) to investigate the difference between genome alignment followed by hard-threshold counting and transcriptome alignment followed by probabilistic counting.

Despite the very different approaches to obtaining the count estimates for each gene, the pairwise Pearson correlation coefficients across all the libraries (Fig. S1) were surprisingly similar, with respective differences for each of the 210 pairwise correlation coefficients within 0.03 for both “RAW-Bowtie-RSEM minus RAW-BWA-HTSeq” and “QC-Bowtie-RSEM minus QC-BWA-HTSeq” comparisons. To detect a trend in interlibrary correlation, values across the 4 combinations of read pre-processing and alignment / counting pipelines, we counted for each combination - the numbers of times a particular method combination had the maximum correlation value across all four pipelines. Fig.3 shows that the RAW-Bowtie-RSEM pipeline was the overall winner, with 94 maximal values out of 210 comparisons and 12 out of 15 comparisons restricted to biological replicates. If we exclude the read trimming factor, the Bowtie-RSEM pipeline shows overall better performance as compared to BWA-HTSeq by having 130 winners in 210 comparisons (p=0.00034) with the overall pool of libraries and 13 winners in 15 comparisons (p=0.0037) between biological replicates. The effect of read pre-processing was opposite for the two alignment / counting methods: while it improved the interlibrary correlations for BWA-HTSeq, it affected the RSEM results in exactly the opposite way (Fig.3). This is not surprising, considering the nature of the methods: while BWA-HTSeq takes the supplied sequence information as is and therefore heavily depends on the quality of the base calls, RSEM incorporates nucleotide-level quality scores in its model ^16^ which allows the model to leverage the information in the lower-quality 3’-tail of the sequencing reads to construct better alignments. This shows the robustness of RSEM’s model even with a bacterial genome, where the complications of alternative splicing and high share of non-coding genomic DNA (major issues addressed by RSEM) are not applicable.

**Fig. 3.**
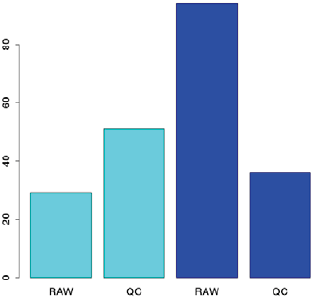
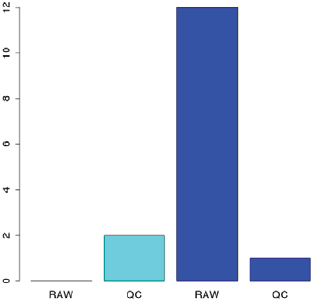
Number of pairwise comparisons a particular combination of read preprocessing and alignment / counting method resulted in the maximal value of Pearson correlation between the genome-wide vectors of counts. **A**, winning cases out of all the 210 pairs, irrespective to experimental conditions; **B**, winning cases out of 15 interreplicate comparisons. Cyan, BWA- HTSeq pipeline; blue, Bowtie-RSEM pipeline.

### Differential expression calling

#### Genome-wide profiles of significance values of DE calling

Fig. S2 shows distributions of gene-level statistics for DE tests performed with 3 methods (DESeq, EBSeq, edgeR and voom / limma). The overall profiles of significance level assignments are found to be dramatically method-dependent. Based on biological reasoning, when one may expect genes to be either differentially expressed between the conditions or not, the ideal-world genome-wide profile of DE significance value is expected to be bimodal with peaks near the extreme (0 or 1) values, with low density in the intermediate zone that represents “class assignment hesitation” of an algorithm. Still, lack of discrimination against the “hesitation zone” was very obvious for 3 out of 4 algorithms (Fig. S2 A, C, D). On the contrary, EBSeq (Fig. S2B) has demonstrated a trend of “hesitation zone avoidance”. While the latter observation is encouraging, the distribution profile alone cannot be taken as an evidence for a better biological relevance of EBSeq-generated output without other, complementary evaluation strategies.

#### Individual library-level expression overlaps between the conditions

The majority of the transcriptomic experiments (both microarray and RNA-Seq) have a low number of biological replicates, with 3 or even 2 replicates dominating the overall pool of publically available repositories such as GEO^17^. Given that fact, which represents the *de facto* limited information on true data variability for any given gene, a biologist conducting the experiment needs to stay on conservative side and focus on genes that show a consistent trend of differential expression between the conditions at the individual replicate level. With 2-3 replicate experiments, having a replicate which shows a trend of expression change that is opposite to the mean trend between the two conditions is intuitively unreliable, and such genes would not be chosen for follow-up by an experimenter. To formalize this selection, we introduced a “critical coefficient” (crt) filter which is the ratio of the minimal expression level found across all the replicates the belong to the condition with higher mean expression level to the maximal expression level found across all the replicates the belong to the condition with lower mean expression level. Critical coefficient below 1 indicates the presence of unfavorable overlap of replicate-level expression values between the two conditions. We have actually used this filter in a number of earlier microarray studies^18,19,20,21,22^, which helped to improve the sensitivity –specificity balance and discover more consistent biological stories, according to our in-house comparisons with alternative methods. The formal statistical properties of the critical coefficient will be reported elsewhere.

While the expectation of replicate-level consistency of expression changes in low-replicate transcriptomic experiment stays in place after transitioning the field from microarrays to RNA-Seq, the relevance of the critical coefficient in RNA-Seq world is not obvious, given the statistical sophistication implemented in the today’s RNA-Seq DE calling methods. Fig.4 shows critical coefficients plotted for the genes called differentially expressed (i.e. with FDR < 0.05) in SynH+LT vs. SynH at timepoint T2 by 4 methods. Surprisingly, 2 out of 4 methods (Fig.4 A, C) demonstrate noticeable presence of genes with overlapping library-level expression values between the conditions (log10(crt) < 0). The phenomenon included pronounced cases like gene “b4354” (Fig. S3) which had extremely low (<0.1) value of critical coefficient indicating dramatic overlap of the expression values between the conditions. While 3 out of 4 methods associated high FDR values with this gene (EBSeq: 0.87; DESeq: 0.60; voom/limma: 0.97), edgeR selected this gene as highly differentially expressed (FDR = 0.0000082). This fact should not be treated as an evidence for a particular inaccuracy of edgeR but rather as an evidence for meaningfulness of critical coefficient filtering on top of modern RNA-Seq differential expression calling methods. It should be noted that the genes selected as differentially expressed by voom / limma showed higher critical coefficient values overall, with a clear trend of those values to follow the statistical significance of the DE call; expression values for all the genes that were called differentially expressed by voom / limma at lower (< 0.005) FDR values were very well (at least 1.8 times) separated between the conditions at the level of individual libraries, while other methods assigned this strict level of statistical significance to numerous genes with weakly separated or even overlapping library-level values (Fig.4). This agreement between voom / limma output and biological common sense expectations suggests that this recent adaptation of a well-developed microarray analysis framework to RNA-Seq data ^23^ deserves closer attention.

**Fig. 4.**
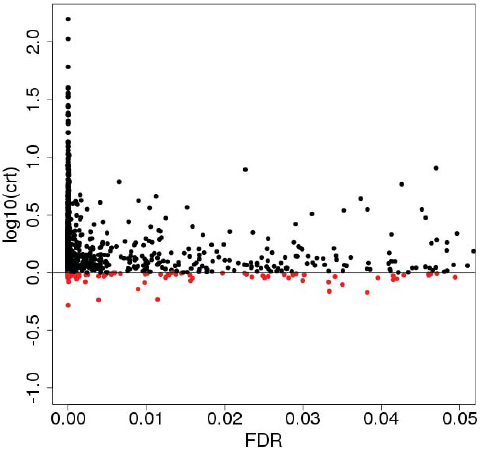
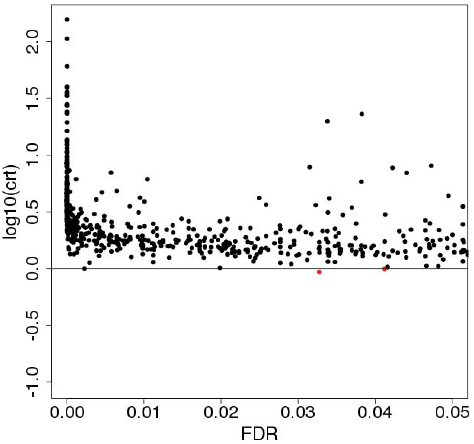
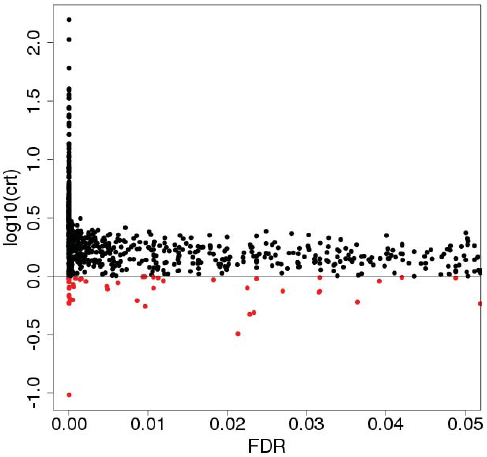
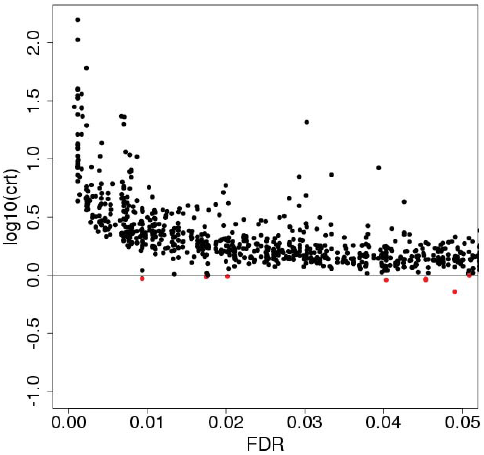
Critical Coefficients computed for genes that are called differentially expressed (FDR < 0.05) by four DE calling methods. For EBSeq, posterior Probability of Equal Expression was used as a conservative estimate of FDR. **A**, EBSeq; **B**, DESeq; **C**, edgeR; **D**, voom / limma. Red, genes with critical coefficient values below 1 (corresponds to 0 in the log-scale applied).

#### Consistency with microarray results

The same experimental design was used in the earlier study ^14^ employing microarray platform. We have used this publically available dataset to assess the robustness of differential expression results across the platforms. Since the ground truth is not available and both microarray and RNA-Seq platforms possess a mixture of features of positive and negative connotation (e.g. proven technology with established analytical toolset but low dynamic range and cross-hybridization problems vs. new technology with potentially higher precision but immature analytical toolset), we applied very neutral / agnostic consistency criterion defined as a ratio between number of genes called DE by both platforms (with same directionality of change) to the size of the pool of genes called DE by any platform, for every combination of the pipeline parameters and biological factors. The value of this Relative Intersection (RI) statistic varied widely (Fig. S4), from almost no overlap (0.01) to about ½ overlap (0.48) at gene level, with median of 0.19 and mean 0.24. Surprisingly, both read pre-processing and the choice of alignment / counting pipeline had clearly no influence of the interplatform DE result consistency (Fig. S5). Moving downstream the processing pipeline, beyond count values generation, we start to see the influence of the processing options: count normalization method showed a borderline influence on the RI (Fig. S6A), and choice of the method for subsequent DE test (Fig. S6B) had significant effect on this statistic. Still, biological factors such as nature of the toxic medium (ACSH or SynH+LT – Fig. S7A), as well as growth stage (Fig. S7B) had the most pronounced effect on the RI.

Complementing the one-factor-at-a-time evaluation, we also took all the considered factors as parts of a generalized linear model that targets RI as a function of all the factors mentioned above plus the critical coefficient (the effect of the latter was not obvious during one factor at a time evaluation – not shown). The model confirmed the highest significance of biological factors, followed by DE calling choice and insignificance of the low-level processing options (Table 1). Artificially toxic (better controlled) medium resulted in better result correspondence across platforms, the latter decreased at stationary phase of growth when interference of multiple biological responses is observed. EBSeq, followed by edgeR, showed elevated RI values, compared to other 2 DE methods. Surprisingly, application of critical coefficient was neutral in terms of resulting interplatform agreement.

**Table 1.** Generalized Linear Model of the RNA-Seq - microarray DE results concordance (expressed as RI) as a function of the technical and biological factors combined. Levels of factors reported: QCRAW, no read pre-processing; alignRSEM, RSEM alignment / counting pipeline; normTMM, count normalization using TMM; normUPPER, count normalization using upper quartile; compsynHLT, biological comparison of SynH+LT vs. SynH; timeT3, transitional phase of cellular growth; timeT4, stationary phase of cellular growth; DEEBSeq, EBSeq as the DE calling algorithm; DEedgeR, edgeR as the DE calling algorithm; DEvoom, voom / limma as the DE calling algorithm; critTRUE, critical coefficient of 1.15 applied. Bold, variables retained after forward-backward regression and p-value filtering.

| Factor | Coefficient | Std. Error | t value | Pr(>\|t\|) |
| --- | --- | --- | --- | --- |
| QCRAW | 0.001 | 0.005 | 0.295 | 0.768 |
| alignRSEM | -0.007 | 0.005 | -1.332 | 0.183 |
| normTMM | -0.005 | 0.006 | -0.854 | 0.393 |
| normUPPER | -0.017 | 0.006 | -2.753 | 0.006 |
| **compsynHLT** | 0.150 | 0.005 | 29.909 | < 2e-16 |
| **timeT3** | -0.024 | 0.006 | -3.925 | 9.74e-05 |
| **timeT4** | -0.172 | 0.006 | -27.971 | < 2e-16 |
| **DEEBSeq** | 0.069 | 0.007 | 9.677 | < 2e-16 |
| **DEedgeR** | 0.045 | 0.007 | 6.361 | 4.15e-10 |
| DEvoom | 0.005 | 0.007 | 0.772 | 0.440 |
| critTRUE | -0.003 | 0.005 | -0.610 | 0.542 |

Overall, there is a trend of increasing the impact of the technical processing stages from upstream pre-processing to downstream differential expression testing. Keeping in mind the fact that the final result of the transcriptomic experiment is inferred pattern of functionality, evaluating the pipeline on the basis of functional pattern conservation is especially informative.

### Relative impact of processing stages on the derived pattern of functionality

#### Overall model

We used 4 types of gene sets representing different dimensions of cellular functionality: 133 KEGG pathways, 400 species-specific pathways, 172 regulons and 333 transporters, combined with 3 gene set enrichment strategies (see Materials and Methods). For every combination of read pre-processing, count normalization, DE test, direction of expression change, gene set type, gene set enrichment strategy and cellular growth stage, we computed the correlation coefficient between the two (ACSH vs. SynH and SynH+LT vs. SynH) vectors of – log10(GS-FDR) within each gene set type, where GS-FDR is the FDR associated with call of involvement of a particular gene set in the transcriptional response. Hence, for KEGG pathways we computed correlations between 133-mer vectors, for species-specific pathways – between 400-mer vectors, and so forth.

The global overview of relative contribution of various processing stages on the resulting inferred functionality patterns was generated via the following Generalized Linear Model:

~~~
FunCor ~ PreProcess + AlignCount + Normaliz + DEMethod + DEDirection + GeneSetType + FunSearchType + TimePoint
~~~

where 

~~~
FunCor
~~~

 is the Pearson correlation between vector of –log10(GS-FDR) for ACSH vs. SynH comparison and the respective vector for SynH+LT vs. SynH comparison, 

~~~
PreProcess
~~~

 is the read pre-processing option (RAW or QC), 

~~~
AlignCount
~~~

 is the alignment / counting pipeline (BWA-HTSeq or Bowtie-RSEM), 

~~~
Normaliz
~~~

 is the gene length-agnostic count normalization method for subsequent DE test (MED, UPPER, TMM), 

~~~
DEMethod
~~~

 is the algorithm for DE calling (EBSeq, DESEq, edgeR, voom / limma), 

~~~
DEDirection
~~~

 is the direction of the expression changes (up or down), 

~~~
GeneSetType
~~~

 is the type of gene set (KEGG, sPW, TF, Transporters), 

~~~
FunSearchType
~~~

 is the enrichment testing strategy (cluster, Fisher, Fold), 

~~~
TimePoint
~~~

 is the growth stage of the cells (T2, T3, T4).

The model summary is shown in Table 2. The contribution of various processing stages varies dramatically, with a clear trend of dramatically higher impact of biological factors (direction of the expression change and cell growth stage, absolute values of model coefficients 0.28 and 0.25) over technical factors (maximal model coefficient 0.13) and increasing the impact of the stage on preservation of the resulting signature of functionality towards the downstream analysis stages: the pre-processing and alignment / counting stages had no detectable influence of the functional signature preservation at all (“PreProcessRAW” and “AlignCountRSEM” lines in the Table 1), normalization, DE testing options and the choice of gene set type had significant influences, however of low magnitude (model coefficients within 0.08), and the choice of functional overrepresentation strategy had the highest impact among all the technical factors.

**Table 2.** Generalized Linear Model of the correlation between vectors of functionality as a function of the technical and biological factors combined. Levels of factors reported: DEDirectionUP, genes upregulated in toxic media; GeneSetTypesPW, species-specific pathways as gene set type; GeneSetTypeTF, regulons as gene set type; GeneSetTypeTransp, Transporters as gene set type; FunSearchTypepiano, Fisher’s gene-level p-values summarization as enrichment test; FunSearchTypepianoFOLD, gene-level summarization of fold changes as enrichment test; PreProcessRAW, nor read pre-processing; AlignCountRSEM, Bowtie-RSEM alignment / counting pipeline; NormalizTMM, TMM count normalization; NormalizUPPER, upper quartile count normalization; TimePointT3, transitional phase of cellular growth; TimePointT4, stationary phase of cellular growth; DEMethodEBSeq, EBSeq as the DE calling algorithm; DEMethodedgeR, edgeR as the DE calling algorithm; DEMethodvoom, voom / limma as the DE calling algorithm. Bold, variables retained after forward-backward regression and p-value filtering.

| Factor | Coefficient | Std. Error | t value | Pr(>\|t\|) |
| --- | --- | --- | --- | --- |
| **DEDirectionUP** | -0.280 | 0.008 | -36.572 | < 2e-16 |
| **GeneSetTypesPW** | -0.039 | 0.011 | -3.657 | 0.00026 |
| **GeneSetTypeTF** | 0.087 | 0.011 | 8.084 | 9.04e-16 |
| **GeneSetTypeTransp** | 0.078 | 0.011 | 7.204 | 7.42e-13 |
| **FunSearchTypepiano** | -0.023 | 0.009 | -2.480 | 0.0132 |
| **FunSearchTypepianoFOLD** | 0.130 | 0.009 | 13.880 | < 2e-16 |
| PreProcessRAW | 0.008 | 0.008 | 1.035 | 0.301 |
| AlignCountRSEM | -0.001 | 0.008 | -0.152 | 0.87914 |
| **NormalizTMM** | -0.077 | 0.009 | -8.577 | < 2e-16 |
| **NormalizUPPER** | -0.015 | 0.010 | -1.525 | 0.127 |
| **TimePointT3** | -0.073 | 0.009 | -8.078 | 9.50e-16 |
| **TimePointT4** | -0.249 | 0.010 | -25.545 | < 2e-16 |
| **DEMethodEBSeq** | 0.009 | 0.010 | 0.819 | 0.413 |
| **DEMethodedgeR** | 0.083 | 0.011 | 7.766 | 1.12e-14 |
| **DEMethodvoom** | 0.047 | 0.011 | 4.242 | 2.29e-05 |

The two most pronounced effects detected – expression change directionality and cell growth stage – are illustrated of Fig.5 and Fig.6, respectively. Fig.5 shows that the conservation of the inferred functional signature is dramatically higher in the case of downregulated genes, where more than half of the cases have Pearson correlation above 0.5. On the contrary, for upregulated genes we observe not only an overall left-shift of the distribution but also a substantial Gaussian component with peak around 0, representing cases of absence of any concordance between cellular responses to the two toxic media. Similar influence of the cell growth stage may be better illustrated when conditioned by the directionality of expression change (Fig.6), where the gradual divergence between the two toxic media from T2 through T3 to T4 is clearly visible (Fig.6A), with main modes of the correlation values distribution for T2 being above 0.5 and for T4 around 0. With the direction of changes that is favorable for functional signature conservation, i.e. downregulation, all the 3 growth stages have the main peak of the correlation values distributions above 0.5, however the stationary stage (T4) still has a substantial tail at very low correlation values, with significant presence in the anti-correlation zone (Fig.6B).

**Fig. 5.**
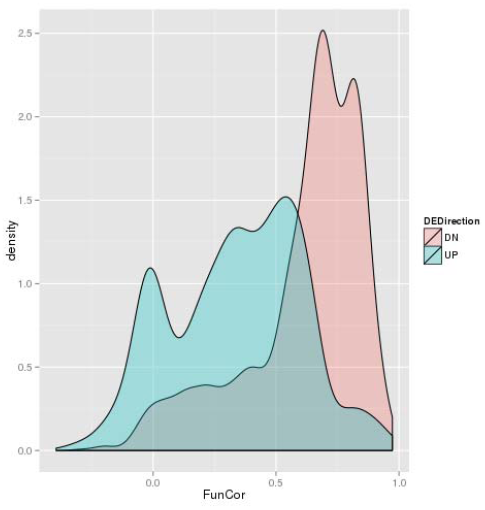
Distributions of correlation coefficients between vectors of -log10(GS-FDR) values computed for ACSH vs. SynH and SynH+LT vs. SynH comparisons for up- and downregulated genes.

**Fig. 6.**
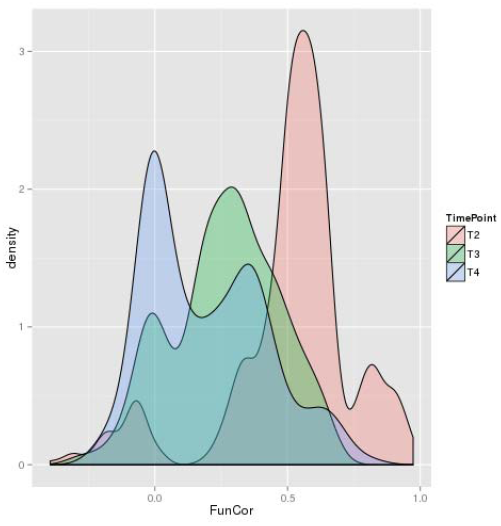
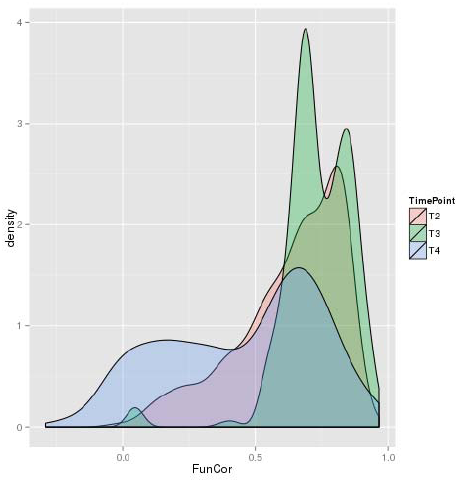
Distributions of correlation coefficients between vectors of -log10(GS-FDR) values computed for ACSH vs. SynH and SynH+LT vs. SynH comparisons for different cell growth stages, with datasets restricted to upregulated **(A)** and downregulated **(B)** genes.

#### Sub-models conditioned by biological factors

While the overall impact distribution between biological and technical factors is clear, one cannot exclude the possibility of masking more subtle technical influences by the overwhelming effect of biological factors. To reveal possible conditional dependences of the impact of technical factors, we constructed the four models with restriction of the data to a particular combination of the most (or the least) favorable directionality of expression change (i.e. downregulation or upregulation) and the most / the least favorable growth stage (i.e. T2 or T4).

TableS1 shows the structures of those models. A key observation there is association of geneset type significance with the direction of the expression change. For example, regulons are positively associated with preservation of functional signatures between the 2 media in the case of upregulated genes (TableS1AC), and no such association is visible for downregulated genes (TableS1BD). The phenomenon is even more pronounced with transporters, when a positive effect in the case of upregulated, and negative effect for downregulated genes is observed (the latter, however, is below statistical significance for growth stage T2). As to the choice of the enrichment method, summarization of fold changes has very positive effect for downregulated genes (TableS1BD), which may be attributed to generally more pronounced effect of downregulation than upregulation in the toxic media. The latter method was also beneficial (however, with less magnitude) for upregulated genes at T2 (TableS1A).

A noticeable phenomenon is the often-seen negative effect of TMM normalization on the functional signature conservation, visible in both the overall model (Table 2) and in condition-restricted models (TableS1ACD). Visualizing the distributions of functional correlation values for upregulated genes at growth stage T2, for example (Fig.7) revealed that while all the 3 normalization methods have the main mode of distribution density above Pearson correlation of 0.5 and no simple scaling method (UPPER or MED) results in functional correlations below 0.12, TMM has a large low-end tail heavily exposed to the anticorrelation zone (Fig.7).

**Fig. 7.**
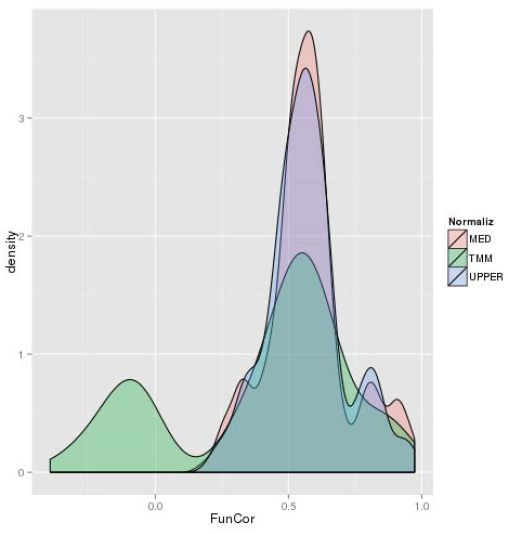
Distributions of correlation coefficients between vectors of -log10(GS-FDR) values computed for ACSH vs. SynH and SynH+LT vs. SynH comparisons for different count normalization methods, with datasets restricted to upregulated genes at exponential (T2) growth stage.

The above observations suggest that choice of the gene set and even functional enrichment search strategy are deeply merged with a biological story.

## Discussion

In this work, we approached a “bird view” evaluation of all the stages of RNA-Seq analysis pipeline in order to estimate relative contribution of option variations at every stage of the pipeline on the resulting functional signature of the cellular response.

The main observation of this study is the vanishing influence of low-level processing stages, such as read pre-processing and alignment / counting options on the functional signature derived downstream. While not all the combinations of read alignment and counting were checked, the results suggest that this level of detail is not necessary for our purpose; the two alignment / counting pipelines, despite very different nature of the underlying methods, had negligible effect on the inferred functional patterns. Still, analysis of interlibrary correlations within the alignment / counting stage and their combined dependence on upstream read pre-processing has demonstrated the advantage of probabilistic counting method (RSEM) that apparently can take advantage of lower-quality base calls to construct better alignments.

Checking relative agreement between RNA-Seq and microarray DE results on gene level discovered an overwhelming influence of biological factors on the agreement statistic, as compared to the technical parameters of the pipeline. Dependence of interplatform concordance on the magnitude of biological effect was reported recently ^24^. Still, it was somewhat counterintuitive that processing options of different nature affect the interplatform concordance in much lesser extent. Interestingly, the work of ^24^ compared 3 (limma, DESeq, edgeR) out of 4 DE methods applied to our study and found the results to be highly correlated. Our observation of different behavior of those methods in terms of genome-wide significance level distribution and preserving the consistency of the direction of expression changes provides another perspective and may motivate further developments of DE calling algorithms with consideration of biological expectations.

The main approach to method judgment applied here is conservation of functional signatures between 2 biologically related comparisons. While this approach is entirely different from microarray – RNA-Seq agreement evaluation, the bigger picture conclusions from the two were very similar: 1) biological factors have the most influence on the final result, 2) earlier stages of the processing pipeline have less impact on the final result than later stages, even if we vary the definition of the “final result” and consider either gene-level differential expression result across 2 platforms or geneset-level results within the same platform across two biologically related comparisons.

Examples of concordant conclusions from the two complementary types of evaluation include: 1) advantage of simple median normalization which demonstrated both better array intersection and better functional pattern reproducibility, 2) overwhelming importance of biological factors: later time point in the experiment (i.e. the stationary phase of cell growth) showed dramatically less agreement with arrays and dramatically less functional conservation between related comparisons within RNA-Seq platform, 3) negligible effect of low-level processing stages. Concerning the item 2, the stationary phase of bacterial cell growth, also known as conditional senescence ^25^ is characterized by co-existence of dying and living cells and transcriptional responses to nutrient limitations. Those factors complicate the transcriptome profile of stationary-phase bacteria, and our observation of lower interplatform concordance at this stage is in agreement with the report that responses involving specific mechanisms show better interplatform concordance than the complex responses ^24^. Beyond interplatform agreement issue at gene level, we show that the same principle is true when applied to within-platform agreement between biologically related comparisons at the level of inferred functionality signatures.

Occasionally, multiple evaluation procedures may result in more questions than answers. For example, DE detection method EBSeq demonstrated best significance values distribution profile, poor consistency of direction of the DE effect across replicates, best coherence with microarray results and none to a marginal advantage on the functionality signature conservation. This confirms that we are far from providing a unified solution which is optimal for all the possible cases. The idea of needing to adapt the analysis to a particular experimental setup is further confirmed by our results on modeling the impact of the processing factors conditioned by the most influential biological factors (TableS1), from which we see large interconnection of the best-performing analytical options and the biological context.

Our results suggest that RNA-Seq analysis should be extremely biology-aware, and special effort should be devoted to optimizing the last stage of the analysis, i.e. search for the functional patterns that form a unique signature of cellular response to the conditions of interest.

## Materials and Methods

### Datasets

RNA-Seq and microarray datasets used in the previous works^14,15^ are available from GEO (accession number GSE58927). Microarray data was processed exactly as described^14^. For RNA-Seq data processing options, see below.

### Reads pre-processing

The options included using reads as they come from the sequencing facility – referred to as “RAW” reads, and performing read trimming, resulting in “QC” reads. “RAW” reads were already free from adaptor and other artificial sequences and had length of 100 nt. To generate “QC” reads, Trimmomatic software^26^ with the following rules: 1) remove the first 12 bases from 5’end, 2) remove any number of nucleotides from 3’end that have the average quality score < 30 in a 3-nt sliding window, keep the trimmed read if 36 or more nucleotides are left.

### Alignment and counting

The genome alignment followed by hard-threshold counting pipeline (“BWA-HTSeq”) included alignment of reads with BWA version 0.7.9a-r786^27^ and subsequent counting of genomic features’ coverage with HTSeq version 0.6.1p2^28^ using default parameters. The transcriptome alignment followed by probabilistic counting pipeline (“RSEM”) included read alignment to the transcriptome with Bowtie version 0.12.7^29^, followed by counting with RSEM version 1.2.4^16^. For both pipelines, information of the type of strand – specificity of the libraries (generated with dUTP protocol) was supplied as “--stranded=reverse” or “--forward-prob 0” for BWA-HTSeq and RSEM pipelines, respectively. Fig. S8 shows our entire low-level processing pipeline (with read pre-processing and alignment / counting options). The pipeline code is available at https://github.com/scienceforever/GLSeq under GNU General Public License version 3.

### Count normalization

The entire set of libraries was pre-normalized as a pool using one of the 3 methods: “MED” – median normalization from EBSeq^30^ package, “UPPER” – upper quartile (75 percentile) normalization from the same package, or “TMM” – trimmed mean of M values normalization^31^ implemented in edgeR^32^ package.

### Tests for differential expression

The pre-normalized count datasets were used as input to the 4 differential expression testing algorithms: EBSeq^30^, DESeq^33^, edgeR^32^ and voom / limma^23^. The default count normalization routines built in the packages were switched off.

### Geneset enrichment analysis

Four types of gene sets – KEGG pathways, species-specific pathways, regulons and transporters – were used. KEGG gene-pathway assignments were downloaded via Bioconductor’s KEGGREST^34^ package, other genesets were formatted from EcoCyc version 17.0 35 flat files. Cluster-based enrichment test with sets of responsive genes (FDR < 0.05) was performed with goseq package^36^, the two gene-level statistics summarization tests – Fisher’s combined probability test^37^ and summarization of median gene-level fold changes - using *piano* ^38^ Bioconductor package. Statistical significance of the enrichment tests was estimated with 100,000 data permutations.

### Generalized Linear Modeling

Generating of the models was performed in R statistical environment. To refine the condition-specific models (presented in Supplementary tables), stepwise forward-backward regression was applied with Bayesian Information Criterion penalty. The models were additionally cleaned up by removing variables with p > 0.05. The initial models (Table 1 and 2) are presented before stepwise regression and p-value filtering, to demonstrate relative contributions of different processing factors explicitly.

## Acknowledgments

This work was supported by the DOE Great Lakes Bioenergy Research Center (DOE BER Office of Science DE-FC02-07ER64494). Authors are thankful to Dr. Dana Wohlbach for advice on BWA processing options and Prof. Robert Landick for general support of the project.

## Abbreviations

DE: differential expression
ANOVA: analysis of variance
FDR: false discovery rate
FPKM: fragments per kilobase of exon per million mapped fragments

## Suppelementary Figures

**Fig. S1.**
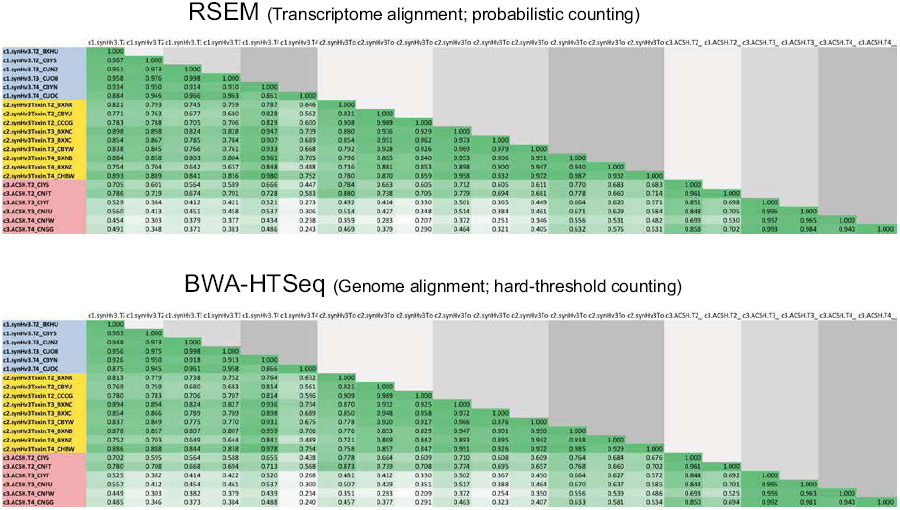
Matrices of interlibrary Pearson correlation coefficients between vectors of counts for all the RNA-Seq libraries in the study. Color code for experimental conditions: blue, control (SynH without lignotoxins) medium; yellow, SynH with lignotoxins; red, ACSH. Shades of grey – growth stages (light to dark: exponential, transitional, and stationary). White-green gradient: color code for Pearson correlation values.

**Fig. S2.**
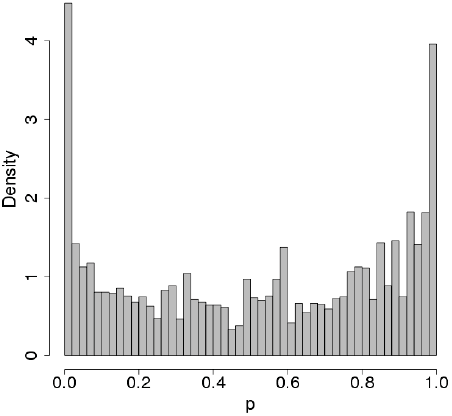
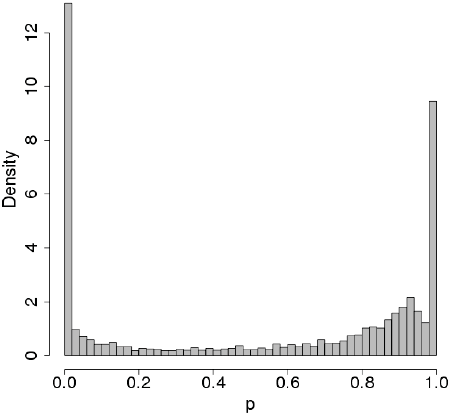
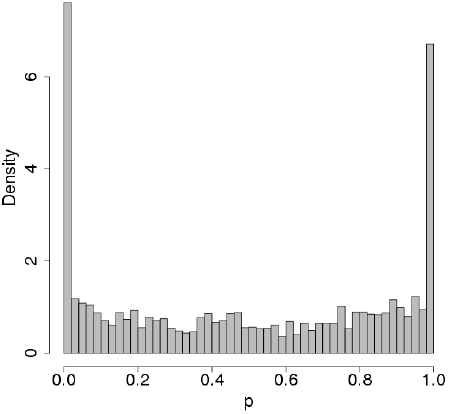
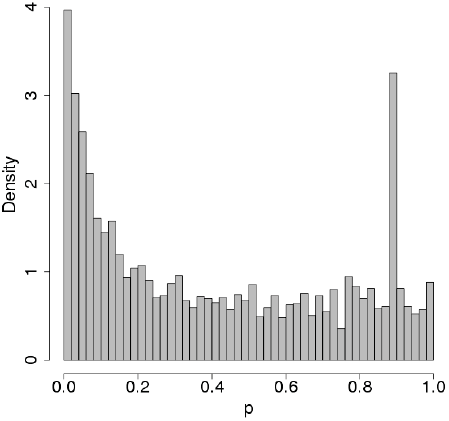
Genome-wide distributions of gene-level significance values for DE calls (A, C, D: Benjamini-Hochberg FDR; B: Posterior Probability of equal expression) for DESeq (A), EBSeq (B), edgeR (C) and voom / limma (D).

**Fig. S3.**
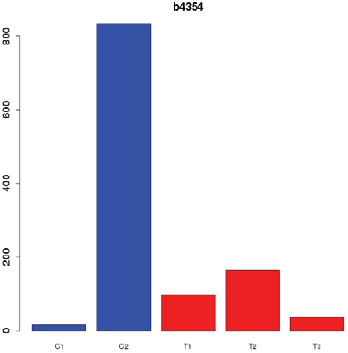
An example of gene with inconsistent library-level expression change between the conditions which was labeled as significantly DE by edgeR. C1-C2, expression levels in SynH; T1-T3, expression levels in SynH+LT. FPKM levels are shown.

**Fig. S4.**
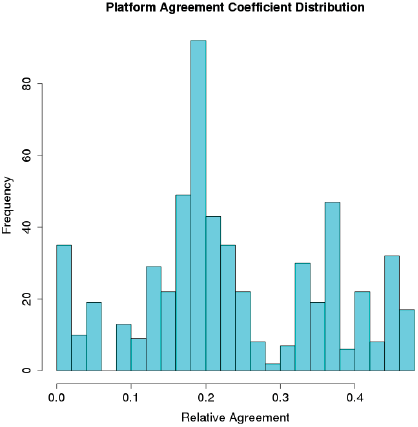
Distribution of Relative Intersection (RI) statistic across 576 comparisons of DE lists between microarray and RNA-Seq results.

**Fig. S5.**
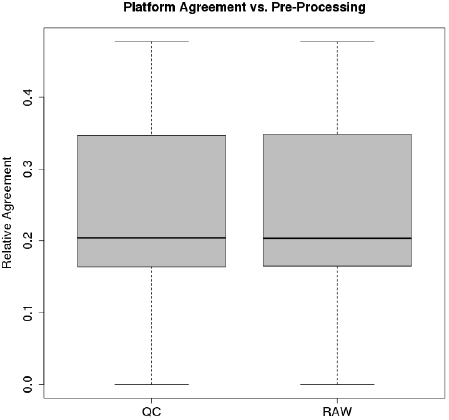
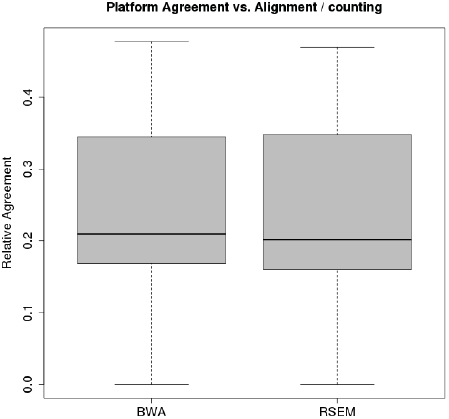
Influence of read pre-processing options **(A)** and the choice of alignment / counting pipeline **(B)** on the RI.

**Fig. S6.**
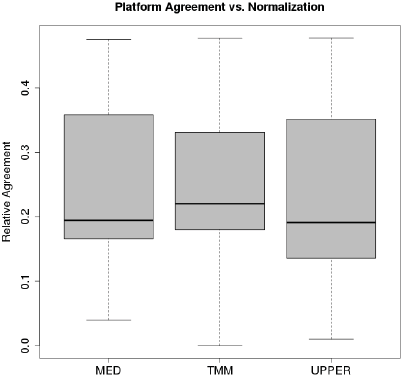
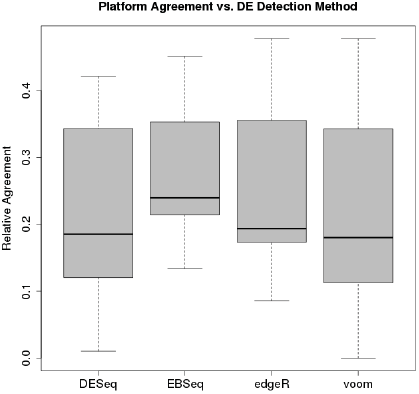
Influence of count normalization **(A)** and DE calling method **(B)** options on the RI.

**Fig. S7.**
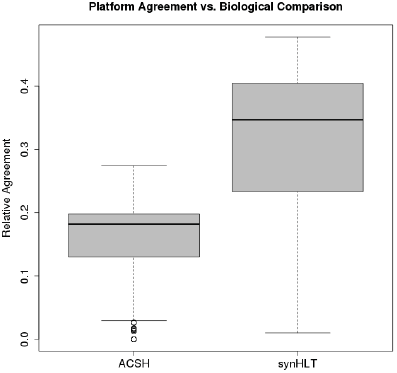
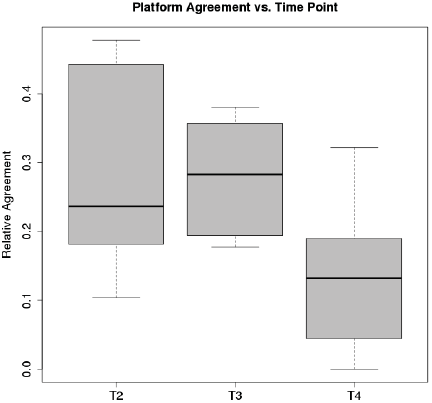
Influence of biological factors: nature of the toxic medium **(A)** and time point **(B)** on the RI.

**Fig. S8.**
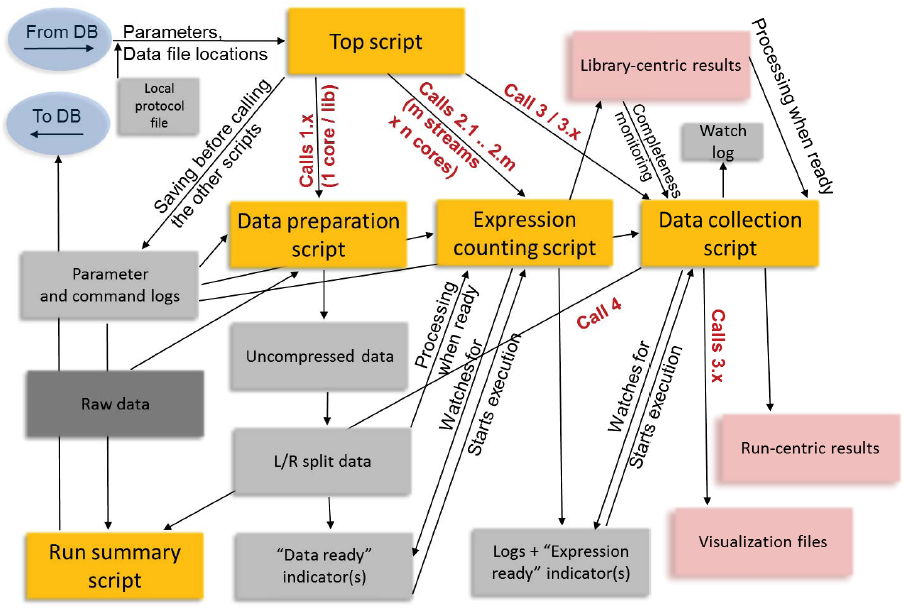
Pipeline for low-level RNA-Seq processing. Yellow, R scripts; dark grey, input data; light grey, intermediate / supplementary files; pink, sets of output files; blue, connections to a database that supplies and stores various run parameters. The top script launches semi-autonomous downstream scripts that watch for the completion of the upstream steps. Both single- and paired-ended libraries and different types of strand-specificity are supported

**Model restricted to upregulated genes at T2 (552 correlations)**

A. Model restricted to upregulated genes at T2 (552 correlations)
| Factor | Coefficient | Std. Error | t value | Pr(>\|t\|) |
| --- | --- | --- | --- | --- |
| GeneSetTypeTransp | 0.295 | 0.023 | 13.1 | 4.85e-34 |
| NormalizTMM | -0.172 | 0.019 | -8.95 | 5.54e-18 |
| DEMethodedgeR | 0.159 | 0.023 | 7.00 | 7.49e-12 |
| DEMethodvoom | 0.154 | 0.023 | 6.66 | 6.71e-11 |
| GeneSetTypeTF | 0.151 | 0.023 | 6.69 | 5.72e-11 |
| FunSearchTypepianoFOLD | 0.093 | 0.020 | 4.74 | 2.71e-06 |
| GeneSetTypesPW | 0.077 | 0.023 | 3.43 | 6.57e-04 |
| DEMethodEBSeq | 0.045 | 0.023 | 1.98 | 4.85e-02 |

**Model restricted to downregulated genes at T2 (549 correlations)**

B. Model restricted to downregulated genes at T2 (549 correlations)
| Factor | Coefficient | Std. Error | t value | Pr(>\|t\|) |
| --- | --- | --- | --- | --- |
| FunSearchTypepianoFOLD | 0.239 | 0.014 | 17.6 | 2.13e-55 |
| GeneSetTypesPW | -0.225 | 0.016 | -14.4 | 6.82e-40 |
| DEMethodEBSeq | -0.037 | 0.016 | -2.33 | 2.02e-02 |

**Model restricted to downregulated genes at T4 (368 correlations)**

C. Model restricted to upregulated genes at T4 (368 correlations)
| Factor | Coefficient | Std. Error | t value | Pr(>\|t\|) |
| --- | --- | --- | --- | --- |
| GeneSetTypeTF | 0.161 | 0.027 | 6.03 | 4.08e-09 |
| GeneSetTypeTransp | 0.159 | 0.028 | 5.64 | 3.42e-08 |
| NormalizTMM | -0.132 | 0.024 | -5.57 | 5.08e-08 |
| DEMethodedgeR | -0.129 | 0.027 | -4.69 | 3.83e-06 |

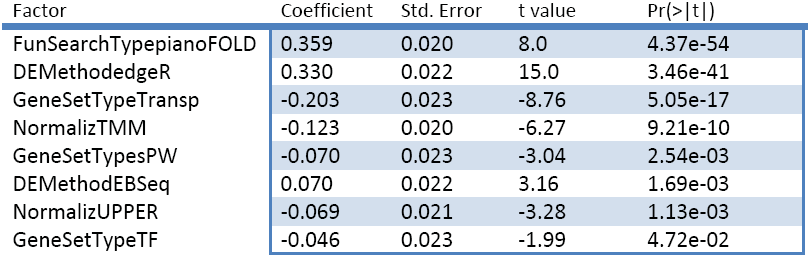
**Model restricted to downregulated genes at T4 (428 correlations)**

C. Model restricted to downregulated genes at T4 (428 correlations)
| Factor | Coefficient | Std. Error | t value | Pr(>\|t\|) |
| --- | --- | --- | --- | --- |
| FunSearchTypepianoFOLD | 0.359 | 0.020 | 8.0 | 4.37e-54 |
| DEMethodedgeR | 0.330 | 0.022 | 15.0 | 3.46e-41 |
| GeneSetTypeTransp | -0.203 | 0.023 | -8.76 | 5.05e-17 |
| NormalizTMM | -0.123 | 0.020 | -6.27 | 9.21e-10 |
| GeneSetTypesPW | -0.070 | 0.023 | -3.04 | 2.54e-03 |
| DEMethodEBSeq | 0.070 | 0.022 | 3.16 | 1.69e-03 |
| NormalizUPPER | -0.069 | 0.021 | -3.28 | 1.13e-03 |
| GeneSetTypeTF | -0.046 | 0.023 | -1.99 | 4.72e-02 |

## References

1. Nagalakshmi U, Wang Z, Waern K, Shou C, Raha D, Gerstein M, Snyder M. The transcriptional landscape of the yeast genome defined by RNA sequencing. Science 2008; 320:1344–9.

2. Blow N. Transcriptomics: The digital generation. Nature 2009; 458:239–42.

3. Mei R, Hubbell E, Bekiranov S, Mittmann M, Christians FC, Shen MM, Lu G, Fang J, Liu WM, Ryder T, et al. Probe selection for high-density oligonucleotide arrays. Proceedings of the National Academy of Sciences of the United States of America 2003; 100:11237–42.

4. Upton GJ, Harrison AP. The detection of blur in affymetrix GeneChips. Statistical applications in genetics and molecular biology 2010; 9:Article37.

5. Irizarry RA, Bolstad BM, Collin F, Cope LM, Hobbs B, Speed TP. Summaries of Affymetrix GeneChip probe level data. Nucleic acids research 2003; 31:e15.

6. Harrison A, Binder H, Buhot A, Burden CJ, Carlon E, Gibas C, Gamble LJ, Halperin A, Hooyberghs J, Kreil DP, et al. Physico-chemical foundations underpinning microarray and next-generation sequencing experiments. Nucleic acids research 2013; 41:2779–96.

7. Mortazavi A, Williams BA, McCue K, Schaeffer L, Wold B. Mapping and quantifying mammalian transcriptomes by RNA-Seq. Nature methods 2008; 5:621–8.

8. Labaj PP, Leparc GG, Linggi BE, Markillie LM, Wiley HS, Kreil DP. Characterization and improvement of RNA-Seq precision in quantitative transcript expression profiling. Bioinformatics 2011; 27:i383–91.

9. Schwartz S, Oren R, Ast G. Detection and removal of biases in the analysis of next-generation sequencing reads. PloS one 2011; 6:e16685.

10. Liu F, Jenssen TK, Trimarchi J, Punzo C, Cepko CL, Ohno-Machado L, Hovig E, Kuo WP. Comparison of hybridization-based and sequencing-based gene expression technologies on biological replicates. BMC genomics 2007; 8:153.

11. Sultan M, Schulz MH, Richard H, Magen A, Klingenhoff A, Scherf M, Seifert M, Borodina T, Soldatov A, Parkhomchuk D, et al. A global view of gene activity and alternative splicing by deep sequencing of the human transcriptome. Science 2008; 321:956–60.

12. Malone JH, Oliver B. Microarrays, deep sequencing and the true measure of the transcriptome. BMC biology 2011; 9:34.

13. Shendure J. The beginning of the end for microarrays? Nature methods 2008; 5:585–7.

14. Schwalbach MS, Keating DH, Tremaine M, Marner WD, Zhang Y, Bothfeld W, Higbee A, Grass JA, Cotten C, Reed JL, et al. Complex physiology and compound stress responses during fermentation of alkali-pretreated corn stover hydrolysate by an Escherichia coli ethanologen. Applied and environmental microbiology 2012; 78:3442–57.

15. Keating DH, Zhang Y, Ong IM, McIlwain S, Morales EH, Grass JA, Tremaine M, Bothfeld W, Higbee A, Ulbrich A, et al. Aromatic inhibitors derived from ammonia-pretreated lignocellulose hinder bacterial ethanologenesis by activating regulatory circuits controlling inhibitor efflux and detoxification. Frontiers in microbiology 2014; 5:402.

16. Li B, Dewey CN. RSEM: accurate transcript quantification from RNA-Seq data with or without a reference genome. BMC bioinformatics 2011; 12:323.

17. Edgar R, Domrachev M, Lash AE. Gene Expression Omnibus: NCBI gene expression and hybridization array data repository. Nucleic acids research 2002; 30:207–10.

18. Braatsch S, Moskvin OV, Klug G, Gomelsky M. Responses of the Rhodobacter sphaeroides transcriptome to blue light under semiaerobic conditions. Journal of bacteriology 2004; 186:7726–35.

19. Moskvin OV, Gomelsky L, Gomelsky M. Transcriptome analysis of the Rhodobacter sphaeroides PpsR regulon: PpsR as a master regulator of photosystem development. Journal of bacteriology 2005; 187:2148–56.

20. Zeller T, Moskvin OV, Li K, Klug G, Gomelsky M. Transcriptome and physiological responses to hydrogen peroxide of the facultatively phototrophic bacterium Rhodobacter sphaeroides. Journal of bacteriology 2005; 187:7232–42.

21. Moskvin OV, Kaplan S, Gilles-Gonzalez MA, Gomelsky M. Novel heme-based oxygen sensor with a revealing evolutionary history. The Journal of biological chemistry 2007; 282:28740–8.

22. Zeller T, Mraheil MA, Moskvin OV, Li K, Gomelsky M, Klug G. Regulation of hydrogen peroxide-dependent gene expression in Rhodobacter sphaeroides: regulatory functions of OxyR. Journal of bacteriology 2007; 189:3784–92.

23. Law CW, Chen Y, Shi W, Smyth GK. voom: precision weights unlock linear model analysis tools for RNA-seq read counts. Genome biology 2014; 15:R29.

24. Wang C, Gong B, Bushel PR, Thierry-Mieg J, Thierry-Mieg D, Xu J, Fang H, Hong H, Shen J, Su Z, et al. The concordance between RNA-seq and microarray data depends on chemical treatment and transcript abundance. Nature biotechnology 2014; 32:926–32.

25. Fredriksson A, Nystrom T. Conditional and replicative senescence in Escherichia coli. Current opinion in microbiology 2006; 9:612–8.

26. Bolger AM, Lohse M, Usadel B. Trimmomatic: a flexible trimmer for Illumina sequence data. Bioinformatics 2014; 30:2114–20.

27. Li H, Durbin R. Fast and accurate short read alignment with Burrows-Wheeler transform. Bioinformatics 2009; 25:1754–60.

28. Anders SP, P.T.; Huber, W. HTSeq - A Python framework to work with high-throughput sequencing data. BioRxiv 2014.

29. Langmead B, Trapnell C, Pop M, Salzberg SL. Ultrafast and memory-efficient alignment of short DNA sequences to the human genome. Genome biology 2009; 10:R25.

30. Leng N, Dawson JA, Thomson JA, Ruotti V, Rissman AI, Smits BM, Haag JD, Gould MN, Stewart RM, Kendziorski C. EBSeq: an empirical Bayes hierarchical model for inference in RNA-seq experiments. Bioinformatics 2013; 29:1035–43.

31. Robinson MD, Oshlack A. A scaling normalization method for differential expression analysis of RNA-seq data. Genome biology 2010; 11:R25.

32. Robinson MD, McCarthy DJ, Smyth GK. edgeR: a Bioconductor package for differential expression analysis of digital gene expression data. Bioinformatics 2010; 26:139–40.

33. Anders S, Huber W. Differential expression analysis for sequence count data. Genome biology 2010; 11:R106.

34. Tenenbaum D. KEGGREST: Client-side REST access to KEGG. R package version 1.4.1. 2013.

35. Keseler IM, Mackie A, Peralta-Gil M, Santos-Zavaleta A, Gama-Castro S, Bonavides-Martinez C, Fulcher C, Huerta AM, Kothari A, Krummenacker M, et al. EcoCyc: fusing model organism databases with systems biology. Nucleic acids research 2013; 41:D605-12.

36. Young MD, Wakefield MJ, Smyth GK, Oshlack A. Gene ontology analysis for RNA-seq: accounting for selection bias. Genome biology 2010; 11:R14.

37. Fisher RA. Statistical methods for research workers. Edinburgh,: Oliver & Boyd, 1932.

38. Varemo L, Nielsen J, Nookaew I. Enriching the gene set analysis of genome-wide data by incorporating directionality of gene expression and combining statistical hypotheses and methods. Nucleic acids research 2013; 41:4378–91.

